# Diagnosis and early prediction of lymphoma using high-throughput clonality analysis of bovine leukemia virus-infected cells

**DOI:** 10.1101/2022.06.06.495041

**Authors:** Tomohiro Okagawa, Honami Shimakura, Satoru Konnai, Masumichi Saito, Takahiro Matsudaira, Naganori Nao, Shinji Yamada, Kenji Murakami, Naoya Maekawa, Shiro Murata, Kazuhiko Ohashi

**Author notes:** Correspondence: Satoru Konnai. These authors contributed equally to this work (Tomohiro Okagawa, Honami Shimakura).

## Abstract

Bovine leukemia virus (BLV), a retrovirus, infects into B cells of ruminants and causes aggressive leukemia or lymphoma in cattle, enzootic bovine leukosis (EBL). Clonal expansion of BLV-infected cells is a promising marker for early detection and diagnosis of EBL. Recently, we developed rapid amplification of the integration site without interference by genomic DNA contamination (RAISING) and CLOVA, a software to analyze clonality. RAISING-CLOVA could assess the risk of adult T-cell leukemia/lymphoma development in human T-cell leukemia virus-I-infected individuals through its clonality analysis. Thus, we herein examined the performance of RAISING-CLOVA for the clonality analysis of BLV-infected cells and conducted a comprehensive clonality analysis by RAISING-CLOVA in EBL and non-EBL cattle. RAISING-CLOVA successfully distinguished EBL from non-EBL cattle with high sensitivity and specificity. A longitudinal clonality analysis in BLV-infected sheep, an EBL model, also confirmed the effectiveness of BLV clonality analysis with RAISING-CLOVA for early detection of EBL development. Therefore, our study emphasizes the usefulness of RAISING-CLOVA as a routine clinical test for monitoring virus-related cancers.

## Introduction

Enzootic bovine leukemia (EBL) is a B-cell lymphoma in cattle caused by infection with bovine leukemia virus (BLV). BLV is a member of the genus *Deltaretrovirus*, subfamily *Orthoretrovirinae*, family *Retroviridae*, and is genetically related to human T cell leukemia virus type 1 (HTLV-1) (1). BLV infects the B cells of ruminants, and its viral RNA is reverse-transcribed into double-stranded DNA, and then integrated into the host genome as a provirus. Most infected cattle are asymptomatic (aleukemic; AL) throughout their lifespan. About 30% of infected cattle develop persistent lymphocytosis (PL), an abnormal proliferation of BLV-infected B cells. EBL occurs in 1–5% of BLV-infected cattle, which present with B-cell malignant lymphoma in lymph nodes and various other organs, leading to a poor prognosis and death (2).

Although BLV has been eradicated in some European countries through the detection and culling of BLV-infected cattle, it is still endemic worldwide (3). The incidence of BLV infection has been increasing during recent decades in Japan; a nationwide survey of cattle conducted from 2009–2011 confirmed BLV infection in 40.9% of dairy cattle and 28.7% of beef cattle (4). The countermeasures against BLV are urgently needed, but there is no commercially available vaccine or therapeutic drug against BLV. Therefore, the herd management by detection, quarantine, and culling of infected cattle is the most effective method of controlling BLV infection. However, this approach is difficult to implement in endemic areas with large numbers of infected cattle. To reduce the economic damage caused by EBL development in such endemic areas, it is effective to detect and cull cattle at high risk of developing EBL in advance.

During the development of EBL, one or few clones of BLV-infected cells undergo clonal expansion. These malignant cells possess identical integration sites of BLV provirus. Therefore, the clonality of proviral integration sites in BLV-infected cells has been considered as a useful marker to diagnose the onset of EBL (5–7). Several methods have been recently developed to analyze transgene integration sites using high-throughput sequencing (HTS), such as ligation-mediated PCR (5, 8), target capture sequencing (6, 9, 10), inverse PCR (11), Linear amplification-mediated PCR (LAM-PCR) (12), and non-restrictive LAM-PCR (13). However, it would be difficult to analyze multiple specimens inexpensively, sensitively, and rapidly by these current methods. Therefore, there is a need to develop a high-throughput method that overcomes the problems of current methods and that can be applied in clinical testing for BLV clonality analysis.

Recently, we developed <u>R</u>apid <u>A</u>mplification of Integration <u>S</u>ites without <u>in</u>terference by <u>g</u>enomic DNA contamination (RAISING) and a clonality analysis software (CLOVA), a highly sensitive, rapid, inexpensive, and high-throughput method to amplify and analyze random integration sites of transgenes in host genomes (14). RAISING and CLOVA were originally developed for the risk assessment of adult T cell leukemia/lymphoma (ATL) development in HTLV-1 carriers (14, 15). The clonality and proviral integration site of HTLV-1-infected cells can be examined by analyzing the sequence of the amplicon from RAISING using the CLOVA software. The clonality value (Cv) of infected cells calculated by RAISING-CLOVA is an effective marker for the prediction of the risk of ATL onset (14).

BLV is considered to utilize similar mechanisms for proviral integration and tumorigenesis as HTLV-1 (9, 16). In our previous study, the proviral integration sites of BLV-infected cells in AL and EBL cattle was successfully amplified using RAISING (14). Sanger sequencing and HTS confirmed the clonal expansion of BLV-infected cells only in the EBL specimen, even though the number of tested specimens was quite limited (*n* = 2) (14). The clonality analysis of BLV-infected cells by RAISING-CLOVA could be an effective method for the diagnosis and prediction of the EBL onset in cattle. Hence, in this study, we performed the comprehensive clonality analysis of BLV-infected cells using RAISING-CLOVA in EBL and non-EBL cattle, and examined its performance in the diagnosis and prediction of the onset of lymphoma in cattle and sheep.

## Materials and methods

### Blood and tumor samples

Peripheral blood of BLV-infected cattle (Holsteins, Blacks, or crossbreds) was collected from dairy and beef farms in Japan between 2017 and 2022. Peripheral blood, lymph nodes, spleen, and various organs were also collected from cattle diagnosed with lymphoma (Holsteins, Blacks, or crossbreds) at Livestock Hygiene Service Centers and Meat Hygiene Inspection Centers in Japan between 2013 and 2022. The blood and tumor samples were kept refrigerated until cell separation. The animal experiments were approved by the Ethics Committee of the Faculty of Veterinary Medicine, Hokkaido University (approval #17-0024). Verbal informed consent was obtained from the owners for the participation of their animals in this study.

Experimental infection of BLV in sheep was conducted at the Research Farm in the Field Science Center, Faculty of Agriculture, Iwate University. Sheep (Corriedale or Suffolk, three months old) were intraperitoneally inoculated with 3.0 × 10^7^ cells of BLV-infected leukocytes isolated from BLV-infected cattle. After the viral challenge, peripheral blood was collected from the BLV-challenged sheep. The procedures were approved by the Iwate University Animal Care and Use Committee (approval no. A201703).

### Cell isolation

Peripheral blood mononuclear cells (PBMCs) were separated from blood samples by density gradient centrifugation using Percoll (GE Healthcare, Chicago, IL, USA). Whole blood was lysed with ACK lysing buffer to separate white blood cells. Separated cells were then washed twice with phosphate buffered saline (PBS, pH 7.4) and filtered through a 40 μm cell strainer (BD Biosciences, San Jose, CA, USA). Tissue specimens were shredded with scissors and filtered using a 40 µm or 100 µm cell strainer (BD Biosciences) to obtain cell suspensions, and washed twice with PBS. Cells were stained with Trypan Blue Stain (Thermo Fisher Scientific, Waltham, MA, USA) and the number of viable cells was measured using a Countess II FL Automated Cell Counter (Thermo Fisher Scientific). Cells were either used immediately or frozen at −80°C until used in experiments.

### Cell lines

Three bovine leukemic cell lines BLV-infected and uninfected cell lines were used in this study: a BLV-infected B-cell line BL3.1 (17) and a BLV-uninfected T cell line BTL26 (18). All cell lines were cultured in RPMI 1640 medium (Sigma-Aldrich, St. Louis, MO, USA) supplemented with 10% heat-inactivated fetal bovine serum (Thermo Fisher Scientific), 100 IU/mL penicillin, 100 μg/mL streptomycin, and 2 mM L-glutamine (Thermo Fisher Scientific) at 37°C and 8% CO_2_.

### Preparation of genomic DNA

Genomic DNA from whole blood samples, PBMCs, and tissue specimens of cattle was extracted using Wizard Genomic DNA Purification Kits (Promega, Madison, WI, USA) or Quick-DNA Miniprep Kits (Zymo Research, Irvine, CA, USA). Genomic DNA from whole blood samples of sheep was extracted using MagDEA Dx SV (Precision System Science, Matsudo, Japan) with a magLEAD 12gC instrument (Precision System Science). Genomic DNA was extracted from mixtures of KU-1 or BL3.1 and BTL26 for a total of 1 × 10^6^ cells using Quick-DNA Miniprep Kits (Zymo Research). The DNA concentrations of the samples were measured by UV absorbance at 260 nm using a NanoDrop 8000 Spectrophotometer (Thermo Fisher Scientific). Agarose gel electrophoresis was performed to check for the degradation of each DNA sample.

### Quantification of BLV proviral load (PVL)

The BLV *pol* gene was measured in the genomic DNA samples of blood and tissue samples of cattle using real-time PCR with a BLV Detection Kit (Takara Bio, Otsu, Japan) with a LightCycler 480 System II (Roche Diagnostics, Mannheim, Germany). A serial dilution series of the positive control DNA for each kit were used to generate calibration curves to determine the copy number of the BLV provirus. Each DNA sample was tested in duplicate. For the blood and tumor samples of sheep, the BLV *tax* gene was measured in the DNA samples using real-time PCR, as described previously (19).

### Diagnosis of BLV infection and EBL

BLV infection in cattle was diagnosed by confirmed by the detection of anti-BLV antibodies using a commercial ELISA kit (JNC, Tokyo, Japan), and by the detection of BLV provirus using real-time PCR. Plasma samples were screened for anti-BLV antibody. Seropositive samples were further tested for the presence of BLV provirus to confirm BLV infection, as described above. Samples that tested positive for the provirus were diagnosed as “BLV-infected.” The number of lymphocytes in blood samples was counted using an automated hematology analyzer (Celltac α; Nihon Kohden, Tokyo, Japan). BLV-infected cattle were classified as aleukemic (AL) or PL based on the lymphocyte counts as follows: AL < 8,000 cells/μL; PL > 8,000 cells/μL.

Blood and tissue samples from cattle with lymphoma were diagnosed as B-cell lymphoma based on immunophenotyping analysis by flow cytometry and/or B-cell clonality analysis by PCR targeting bovine immunoglobulin heavy chain (20) combined with the quantification of BLV provirus by real-time PCR, as described above.

### Amplification of integration sites of BLV provirus by RAISING

The integration sites of BLV provirus were amplified by RAISING as previously described (14), with some modifications. The primers and reagents used in each step are shown in Supplemental Tables 1 and 2. The reaction conditions of each step are shown in Supplemental Table 3. Single-stranded DNA (ssDNA) of the 3’ LTR region of the BLV provirus and the downstream region of the host genome was synthesized from the extracted genomic DNA using the primer BLV-F1 and KOD-Plus-Neo DNA Polymerase (Toyobo, Osaka, Japan). The synthesized ssDNA was purified using a Monarch PCR & DNA Cleanup Kit (New England Biolabs, Ipswich, MA, USA) and was eluted in ultrapure water. Then, poly(A) and poly(G) tails were added at the 3’ end of the purified ssDNA by terminal transferase (New England Biolabs). The double-stranded DNA was then synthesized and amplified by PCR from the poly(AG)-tailed ssDNA using the primers BLV-F2 and NV-oligo-dT-ADP1 and Q5 Hot Start High-Fidelity DNA Polymerase (New England Biolabs). The second PCR was performed using diluted PCR products, the primers BLV-F3 and ADP1-HTS-R1, and KOD-Plus-Neo DNA Polymerase (Toyobo).

### Sequencing and clonality analysis using CLOVA

The products of the second PCR were purified using AMPure XP (Beckman Coulter, Fullerton, CA, USA) and analyzed using Sanger sequencing with a BigDye Terminator v3.1 Cycle Sequencing Kit (Thermo Fisher Scientific) on a 3130Xl or 3730Xl DNA Analyzer (Thermo Fisher Scientific). Clonality analysis of BLV-infected cells was performed using the sequencing signal data of each sample by a CLOVA software (14), an R program that automatically analyzes the clonality value (Cv) of transgene-integrated cells by dividing the average of signal peak area values of 20 nucleotides at 5’ terminal of host genome sequence of the dominant clone by that at 3’ terminal of BLV proviral sequence.

### Statistical analysis

Significant differences among multiple groups were identified using Kruskal-Wallis one-way analysis of variance followed by Dunn’s test. Association of two values were tested using Receiver operating characteristics (ROC) curve analyses were performed to determine the optimal cutoff values, where sensitivity approximates specificity for each risk factor. All statistical tests were performed using GraphPad Prism 6 (GraphPad Software, San Diego, CA, USA). Differences were considered statistically significant at *P* < 0.05.

## Results

### Performance of clonality analysis by RAISING-CLOVA targeting BLV

To examine the detection limit of RAISING targeting BLV, RAISING was performed using a series of DNA samples from mixtures of different percentages of a BLV-infected B-cell line (BL3.1) and a BLV-uninfected strain (BTL26) and the amplified products were analyzed by Sanger sequencing. BL3.1 harbors multiple copies of BLV provirus in the genome (7) and PVL of BL3.1 was 240 copy/100 cells. Thus, DNA samples extracted from specimens including 0.0001% to 100% BL3.1 contain different PVL ranging from 0.0024 to 240 copy/100 cells. The amplified fragments were observed in the electrophoresis and the combined sequences of BLV provirus and host genome were detected in the DNA samples containing 0.012–240 copy/100 cells of provirus (Fig. 1A and Supplemental Fig. 1). The detection limit for RAISING targeting BLV was 0.012 copy/100 cells in PVL, which is comparable with that for RAISING targeting HTLV-1 (0.032% in HTLV-1 PVL) (14). These results suggest that RAISING targeting BLV is sensitive enough to detect proviral insertion sites, even in specimens with low PVL.

**Fig. 1.**
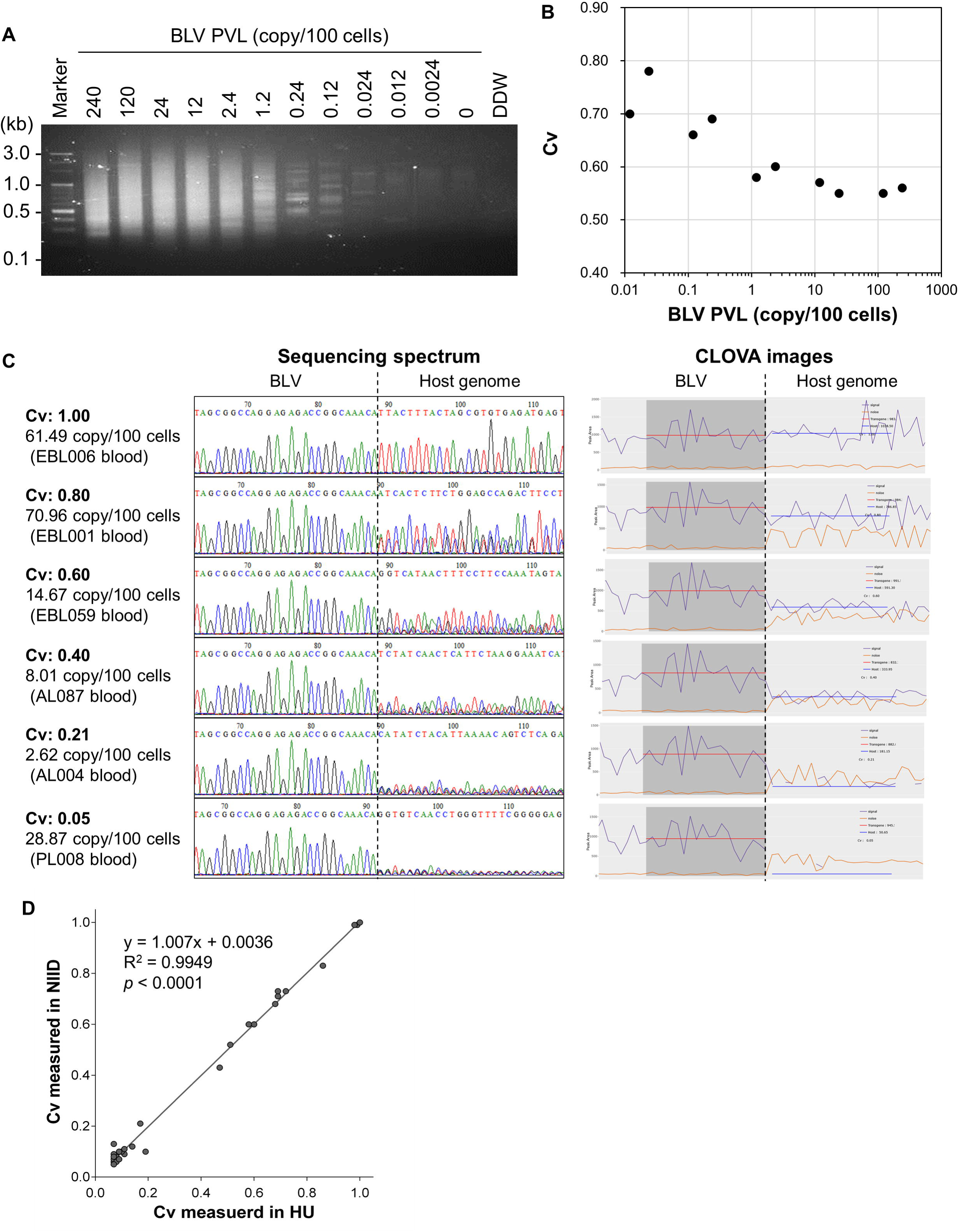
Performance of BLV clonality analysis by RAISING-CLOVA. (A, B) Detection limit of RAISING targeting BLV and sensitivity of its clonality analysis by RAISING-CLOVA. RAISING-CLOVA was performed using genomic DNA from the mixture of the BLV-infected cell line BL3.1 with the BLV-uninfected cell line BTL26. Samples were prepared by mixing BL3.1 with BTL26 in proportions ranging from 100%, 50%, 10%, 5%, 1%, 0.5%, 0.1%, 0.05%, 0.01%, 0.005%, and 0.001% of BL3.1. (A) Products of RAISING using these mixed samples were visualized by electrophoresis on 2% agarose gel. BLV PVL contained in each mixed sample was shown above the photo. (B) The minimum BLV PVL to measure a reliable Cv with RAISING-CLOVA was examined. Sangar sequence spectrum of amplicons of RAISING using a dilution series of BL3.1 were shown in Supplemental Figure 1. (C) Comparison of sequence spectra patterns and signal plots of CLOVA analysis in blood samples of BLV-infected cattle with different clonality values. RAISING-CLOVA was performed using genomic DNA of peripheral blood from EBL and non-EBL cattle. In the plots of CLOVA analysis, signals of dominant clones and the others were shown in purple and orange lines, respectively. The Cv was calculated by dividing the average of signal peak area values of host genome sequence of the dominant clone (red lines) by that of proviral sequence (blue lines). (D) Comparison of Cv analyzed in two different laboratories. RAISING was independently performed at Hokkaido University (HU) and the National Institute of Infectious Diseases (NIID) using the specimens of BLV-infected cattle (*n* = 32) and Cv was determined using CLOVA. Spearman’s rank correlation coefficient was used for statistical analysis.

Our previous study has shown that RAISING-CLOVA targeting HTLV-1 fails to accurately measure Cv in samples with PVL less than 0.5% (14). Here, we examined the sensitivity of clonality analysis by RAISING-CLOVA targeting BLV. The Cv of BL3.1 was 0.56 when measured using the specimen with PVL 240 copy/100 cells (Fig. 1B and Supplemental Fig. 1). The values were comparable (Cv: 0.58) when measured using the specimens with PVL 1.2 copy/100 cells, but it differed by more than 0.1 for specimens with PVL lower than 0.24 copy/100 cells (Fig. 1B and Supplemental Fig. 1). This result indicates that RAISING-CLOVA targeting BLV can accurately measure Cv when the PVL is at least 1.2 copy/100 cells and is feasible enough for the analysis of most BLV-infected specimens.

To further examine the performance and accuracy of the clonality analysis by RAISING-CLOVA, we tested DNA samples obtained from BLV-infected cattle with or without EBL. Representative results of RAISING-CLOVA in blood specimens of BLV-infected cattle with different values of Cv are shown in Fig. 1C. The Cv reflected the intensity of the sequence signal peaks of integration sites at the host genome (Fig. 1C). These results are consistent with our previous results of RAISING-CLOVA using specimens of HTLV-1-infected patients as well as a preliminary analysis using specimens of BLV-infected cattle (14). Furthermore, the clonality analyses of the identical BLV-infected specimens (*n* = 32) in the two different laboratories showed a high interrater agreement in Cv (Fig. 1D). These results indicate that RAISING-CLOVA is very accurate and reproducible method for measuring the clonality of BLV-infected cells.

### Comprehensive clonality analysis of BLV-infected cells in EBL and non-EBL cattle

The clonality analysis by RAISING-CLOVA was performed on peripheral blood samples from AL (*n* = 107), PL (*n* = 79), and EBL cattle (*n* = 101), and tissue samples including tumors from EBL cattle (*n* = 175) collected from farms throughout Japan. In addition, BLV PVL was also measured in these samples, because previous studies proposed that PVL is a candidate marker for EBL diagnosis (21, 22). Representative results of RAISING-CLOVA in EBL and non-EBL blood specimens are shown in Fig. 2A. In an EBL cattle (EBL015), Sanger sequencing analysis detected monoclonal patterns of proviral integration in peripheral blood and tumor specimens and their values of Cv were calculated as 1.00 (Fig. 2A, indicating clonal expansion of BLV-infected cells. In contrast, in non-EBL cattle (AL002 and PL002), the sequencing analysis detected polyclonal patterns of proviral integrations with Cv 0.09 and 0.06, respectively (Fig. 2A). Among the blood samples, EBL showed significantly higher Cv than non-EBL specimens, such as AL and PL (median: 0.63, 0.09, and 0.08, respectively) (Fig. 2B). In contrast, there was no significant difference in the PVL of blood between PL and EBL (median: 35.10 and 25.78 copy/100 cells, respectively), although tumors of EBL showed higher PVL (median: 59.06 copy/100 cells) than blood samples at all disease stages (Fig. 2C). In addition, the Cv of tumors in EBL was higher (median: 0.81) than that of their blood (median: 0.63) (Fig. 2B). Tumors of EBL were observed in a variety of tissues in cattle, including multiple lymph nodes and non-lymphoid tissues such as heart, kidney, uterus, digestive, and respiratory organs. There were no significant differences in the Cv and PVL of tumors from lymphoid and non-lymphoid tissues in EBL (Fig. 2D and E). Furthermore, identical integration sites of BLV provirus were detected by Sangar sequencing among blood and tumor samples from same EBL cattle, even in the blood specimens with low Cv (EBL049 and EBL174) (Supplemental Fig. 2). These results indicate that the Cv of blood samples from RAISING-CLOVA is an effective marker for distinguishing EBL from non-EBL cattle.

**Fig. 2.**
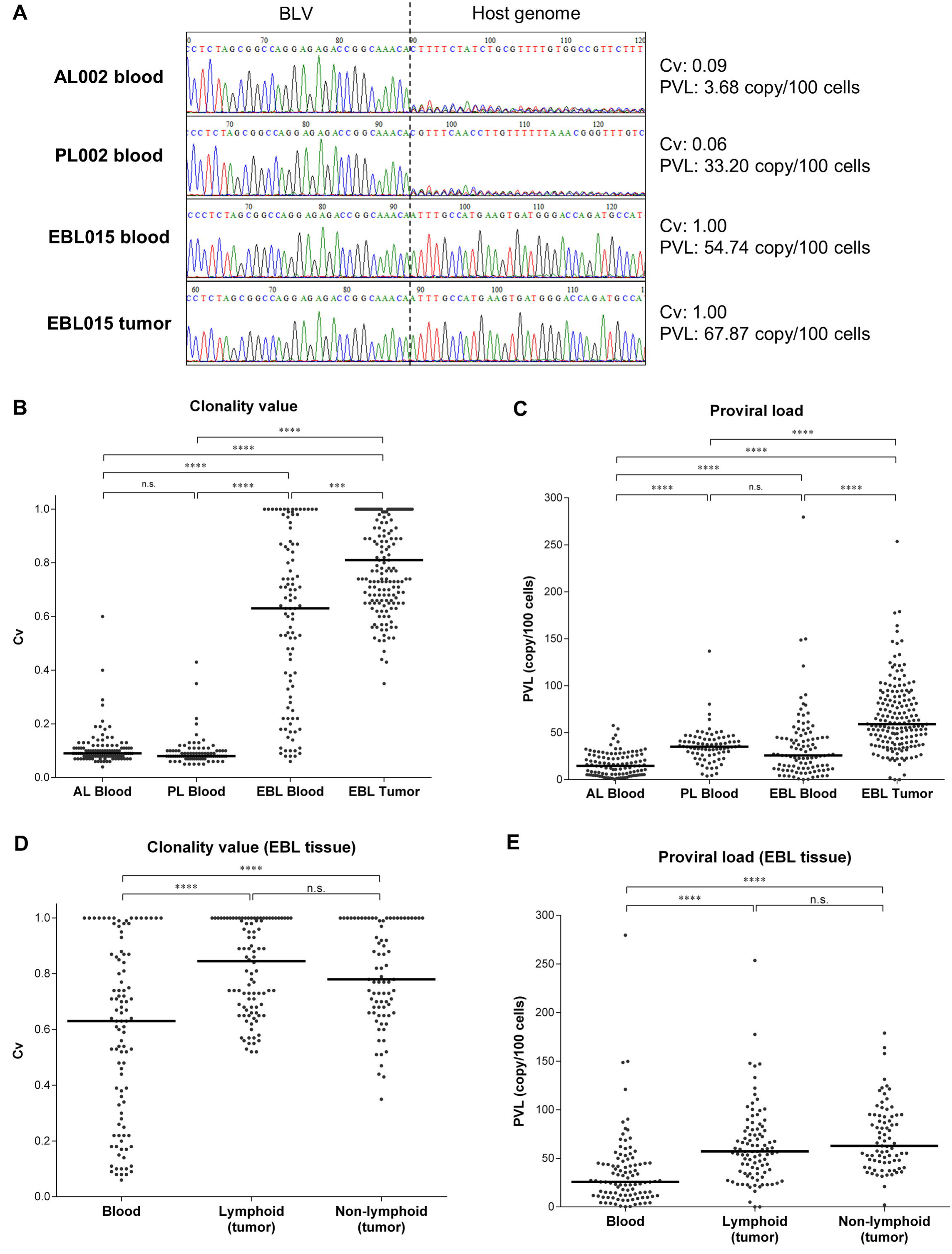
Comprehensive clonality analysis by RAISING-CLOVA targeting BLV using specimens of EBL and non-EBL cattle. (A–E) RAISING-CLOVA was performed using genomic DNA of blood samples of AL (*n* = 107), PL (*n* = 79), and EBL cattle (*n* = 101) and tumor samples of EBL cattle (*n* = 175). (A) Representative Sanger sequence spectrum of blood and tumor samples of BLV-infected cattle (blood samples of AL002, PL002, and EBL015 and a tumor sample from a lymph node of EBL015). (B, C) Cv (B) and BLV PVL (C) was measured in blood samples of AL, PL, and EBL cattle and tumor samples of EBL cattle. (D, E) Cv (D) and BLV PVL (E) among blood and tissue categories in EBL cattle. (B–E) Median values for each group are indicated by black bars. Dunn’s tests were used for statistical analysis. *** *p* < 0.001, **** *p* < 0.0001, n.s., not significant.

**Fig. 3.**
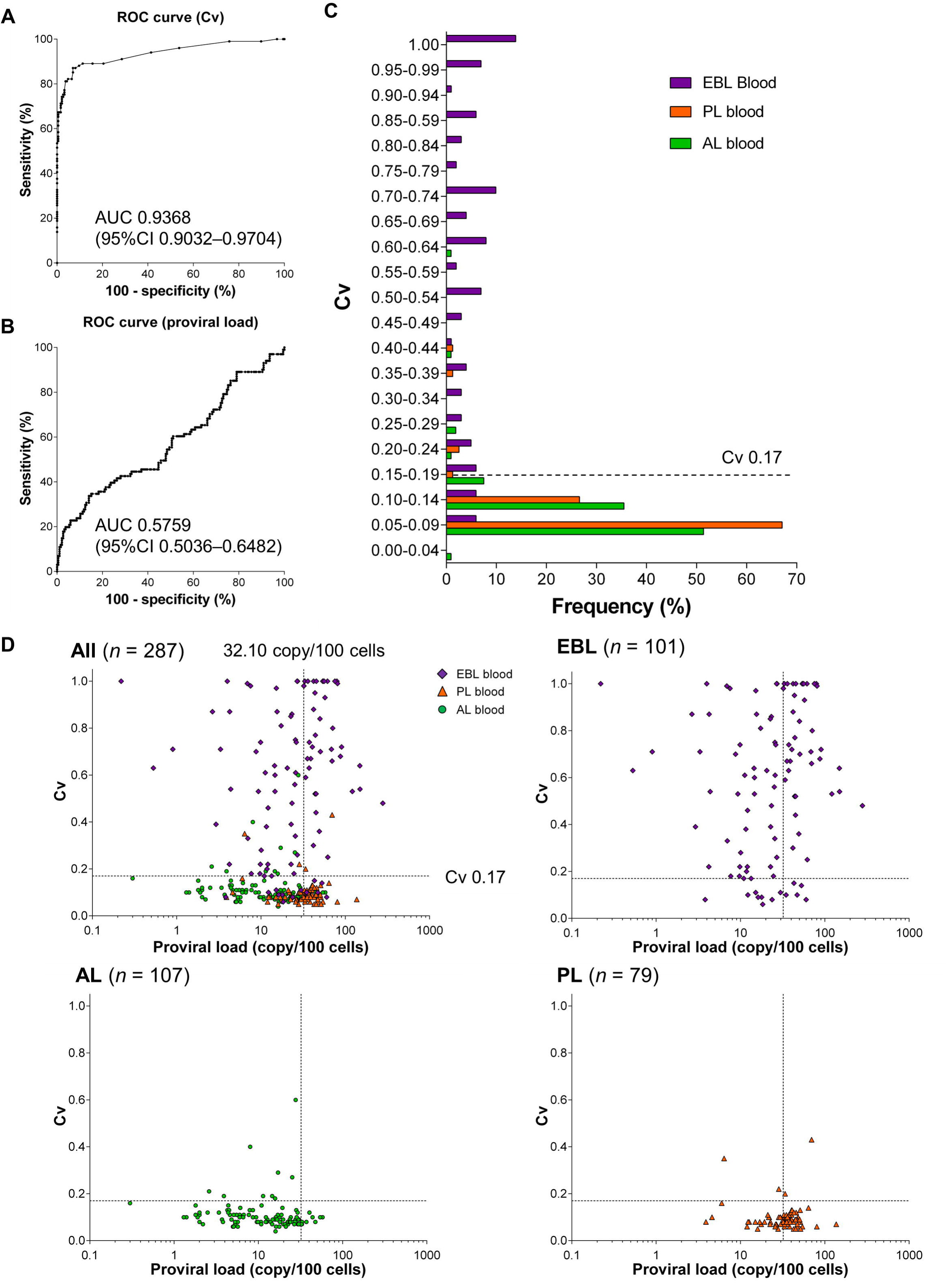
Clinical utility of BLV clonality analysis by RAISING-CLOVA using blood and tumor samples. (A, B) ROC analysis using Cv (A) and PVL (B) of blood samples to distinguish EBL in the blood samples. AUC, area under the curve; CI, confidence interval. (C) Frequencies of blood specimens from EBL (*n* = 101, purple), PL (*n* = 79, orange), and AL cattle (*n* = 107, green) per Cv at intervals of 0.05. A dotted line indicates the proposed cut-off value to classify BLV-infected cattle into EBL. (D) Scatter plots of Cv and BLV PVL of blood samples from EBL (*n* = 101, purple diamond), AL (*n* = 107, green circle), and PL cattle (*n* = 79, orange triangle). Dotted lines indicate proposed cut-off values to classify BLV-infected cattle into EBL.

### Clinical utility of clonality analysis by RAISING-CLOVA targeting BLV

To determine whether the Cv or PVL of BLV-infected cells was valuable for the diagnosis of EBL, we performed ROC analysis and examined the specificity and sensitivity of EBL diagnosis by Cv and PVL in the blood samples. The area under the ROC curve (AUC) for Cv (0.9368) was higher than that for PVL (0.5759) for the blood samples of EBL and non-EBL cattle (Fig. 4A and B). The cutoff value of Cv in blood for EBL diagnosis was 0.17, which could distinguish EBL cases with 87.1% sensitivity and 93.0% specificity (Fig. 4C and D). Out of 101 blood specimens of EBL, the Cv less than 0.17 was detected in 13 specimens (12.9%) (Fig. 4D). These animals showed clonal expansion of malignant cells in tumors, but not in blood. In addition, 13 cases (7.0%) of AL and PL cattle were identified with the Cv > 0.17 in blood (Fig. 4D). Blood samples may also be suitable for follow-up and prognostic studies of non-EBL cattle with intermediate Cv. On the other hand, the cutoff value of PVL in blood was 32.10 copy/100 cells, which was not a suitable marker, because more than half of the samples had lower values than the cutoff (Fig. 4D). Thus, these experiments indicate that Cv is a better diagnostic marker for EBL compared to PVL.

**Fig. 4.**
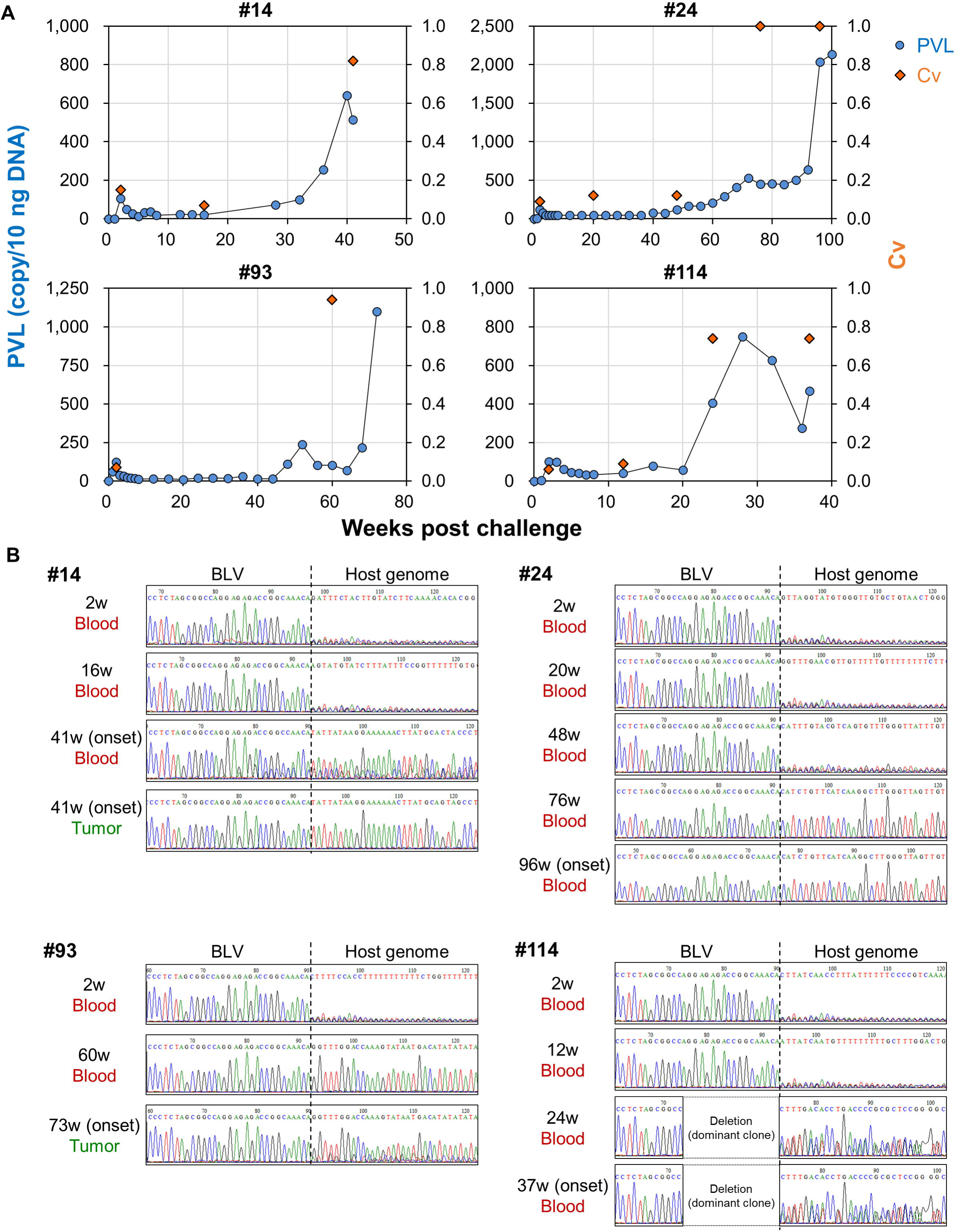
Longitudinal clonality analysis of BLV-infected sheep with lymphoma. (A, B) RAISING-CLOVA was performed using genomic DNA of blood and tumor samples of BLV-challenged sheep (*n* = 4). (A) Kinetics of BLV PVL (blue circle) and Cv (orange diamond) in blood samples of BLV-infected sheep from viral challenge to lymphoma onset. (B) Sequence analysis of the proviral integration sites in blood and/or tumor samples of the challenged sheep.

### Longitudinal clonality analysis of experimental lymphoma model of sheep

Sheep infected with BLV develop lymphoma at higher frequencies after shorter latency periods than cattle (2). To further investigate the usefulness of Cv as a predictive marker for the development of EBL, longitudinal clonality analysis was conducted in an experimental infection model of sheep with BLV. In Cv remained low immediately after infection, but increased before or at the onset of lymphoma (Fig. 5A). In three of the four tested animals (#24, #93, and #114), Cv peaked earlier than PVL in blood (Fig. 5A). Sequence analysis of the integration sites in blood and tumor tissues also revealed that the identical integration sites were detected before and at the onset of lymphoma in all tested sheep (Fig. 5B). These results indicate that clonality analysis by RAISING-CLOVA is a promising method for early prediction of lymphoma onset in BLV infection.

## Discussion

When a retrovirus infects a host, it may persist in the host throughout its lifespan, causing severe disease in some carriers. In the cattle industry, BLV infection is widespread in all parts of the world except Western Europe (3), and causes major economic losses in the production of milk and beef (23, 24). However, there is no effective treatment or vaccination to control BLV infection. In BLV-endemic countries such as Japan and the United States, the control measure of eradicating BLV-infected cattle is completely impractical. Therefore, the development of a novel method for early prediction of tumor development in the carrier stages would contribute to the reduction of economic losses in livestock production. In this study, we applied a novel molecular method called RAISING with the clonality analysis software CLOVA (14) to amplify BLV proviral integration site in host genome and analyze the clonality of BLV-infected cells. We further examined the usefulness of the clonality analysis by RAISING-CLOVA as a method for EBL diagnosis and the early prediction of lymphoma onset.

For BLV clonality analysis, several molecular methods have been developed to analyze proviral integration sites of BLV-infected cells, including ligation-mediated PCR (5), target capture sequencing (6, 9), and inverse PCR (7, 11, 25). Most of these methods requires HTS analysis for the detection of integration sites, but the high cost of the analysis makes it unsuitable for clinical diagnosis with large numbers of samples. Additionally, these current methods raise concerns about the sensitivity and bias of detection, which use restriction enzymes or ultrasound sonication for DNA fragmentation.

RAISING is a highly sensitive, highly accurate, rapid, inexpensive, and high-throughput method to overcome the problems of conventional methods (14). In this study, we analyzed Cv of BLV-infected cells in EBL and non-EBL cattle using RAISING-CLOVA, and found that Cv discriminated between non-malignant and malignant samples successfully but BLV PVL did not. In previous studies, PL and EBL-infected cows are known to have higher provirus levels than AL cows (21, 26). Therefore, proviral levels are considered to be an important marker in EBL diagnosis (Ohno et al., 2015; Kobayashi et al., 2019). However, PVL was not a suitable marker for EBL diagnosis in this study. This inconsistency is presumably because a large number of samples was examined in this study, including PL cows with high PVL. The clonality analysis by RAISING-CLOVA should be conducted with another cohort of clinical samples to confirm the reproducibility of the present analysis.

Among EBL cases analyzed in this study, 13 specimens (12.9%) did not show tumorigenesis in peripheral blood, a finding which is consistent with our previous report (Nishimori et al., 2017). In EBL cattle, lymphocytosis and/or the presence of atypical lymphocytes is reportedly observed in the peripheral blood. but this does not appear to be the case in all cases. However, more detailed EBL typing may be possible by examining the clonality of tumor cells in the blood and tissues of a larger number of EBL cattle. In addition, recent studies suggest that tumor cells of EBL sometimes harbor defective proviruses (6, 11). EBL diagnosis using qPCR targeting provirus would miss those defective cases. In this study, several tumor cells with proviruses deficient in the *pol* gene, which was the target of qPCR in this study, were identified in EBL tumors (data not shown). In BLV-specific forward primers targeting the 3’LTR and its upstream regions, these regions seemed to be less likely to be deleted in the provirus of EBL tumors (6, 11) and could be an optimal target of primers.

In BLV infection, antisense transcripts, called *AS*, were found to be constantly expressed in EBL tumors through promoter activation of the 3’ LTR region (27). A recent study reported that activation of antisense transcription originating from the 3’ LTR forms a chimeric transcript of the *AS* gene and the host driver gene upstream of the provirus, resulting in enhanced transcription of driver genes during BLV infection and following tumorigenesis (9). Therefore, it is important to analyze the integration site of BLV provirus in the bovine genome in order to investigate its contribution to tumorigenesis of EBL. Further studies are warranted to address this issue by HTS analysis of the amplicon of RAISING in non-EBL and EBL cattle.

To further validate the usefulness of clonality analysis by RAISING-CLOVA as a diagnostic method, a comprehensive follow-up study of non-EBL cattle is required in a clinical setting, focusing on carrier animals identified to have higher Cv in the screening test. Previous studies have demonstrated that clonality analysis of HTLV-1-infected patients showed that the Cv of infected cells increased earlier than the increase in peripheral blood provirus levels before the onset of ATL, suggesting the applicability of this method to the risk assessment of ATL (14, 15). In this study, similar results were confirmed in a sheep lymphoma model. Sheep can be a good model for validating the usefulness of this method, because they develops lymphoma at high frequencies and with shorter period compared to cattle (2). We will also test additional sheep samples for further validation. Taken together, further studies of BLV clonality analysis by RAISING-CLOVA will contribute to the establishment of a new control measure to predict disease prognosis of non-EBL cattle and prevent EBL onset by exposing high-risk animals.

## Acknowledgments

We are grateful to all farmers and veterinarians who provided clinical samples used in this study. We thank Enago (http://www.enago.jp) for the English language review.

## Author contributions

TO, SK, MS, NM, SM, and KO designed the work. TO, HS, MS, TM, NN, and SY performed the experiments. MS, TM, NN, SY, and KM provided intellectual input, field samples, laboratory materials, reagents, and/or analytic tools. TO, HS, SK, MS, TM, NN, SY, and KM acquired, analyzed, and interpreted the data. TO and HS wrote the manuscript. SK, MS, SY, KM, NM, SM, and KO revised the manuscript. All authors read and approved the final manuscript.

## Data availability statement

The datasets used and analyzed in this study are available from the corresponding author on reasonable request.

## Additional Information

MS, TM, and NN have a patent pending for materials and techniques described in this paper (application number PCT/JP2020/30907). The other authors declare no competing interests. This work was supported by grants from Ito Memorial Foundation (to SK), the Science and Technology Research Promotion Program for Agriculture, Forestry, Fisheries, and Food Industry, Japan (number 26058B; to SK), the NARO, Bio-oriented Technology Research Advancement Institution (the special scheme project on regional developing strategy; grant 16817557 to SK), and grants-in-aid for Scientific Research (project numbers 19KK0172 and 22H02503 to SK, 19K15993 and 22K15005 to TO, 17H03594 to MS). The funders had no role in study design, data collection and interpretation, or the decision to submit the work for publication.

