## Supplemental Figures 1 and 2, Supplemental Tables 1-3 for "Diagnosis and early prediction of lymphoma using high-throughput clonality analysis of bovine leukemia virus-infected cells"

### Supplemental Figure 1

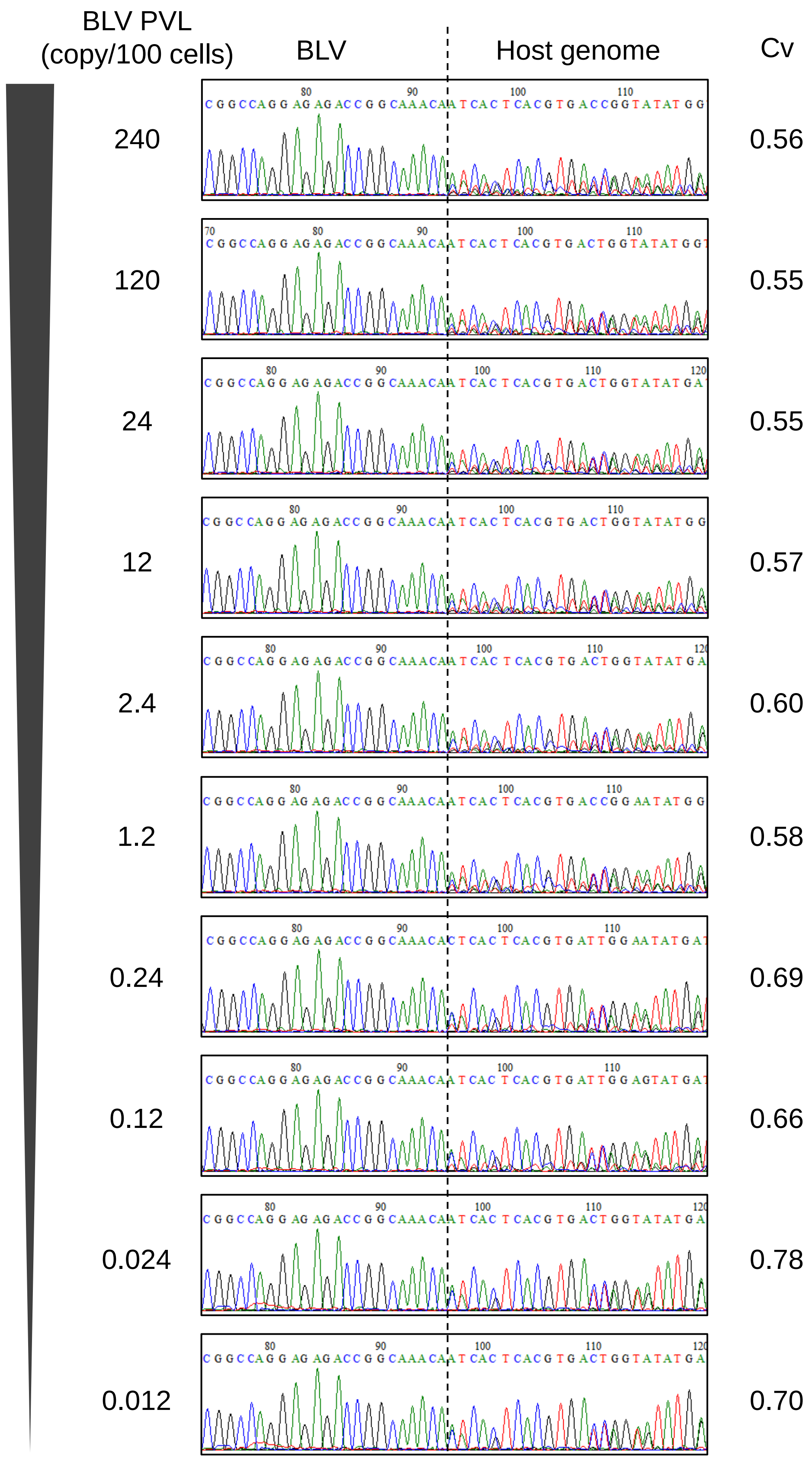


**Supplemental Fig. 1. Sanger sequence spectrum of a dilution series of BLV-infected cell line by RAISING.**

Sangar sequence spectrum of amplicons of RAISING using a dilution series of BLV-infected cell line (BL3.1) were shown with BLV PVL measured by qPCR and Cv analyzed by CLOVA. Dashed line indicates the position of the BLV integration site.

### Supplemental Figure 2

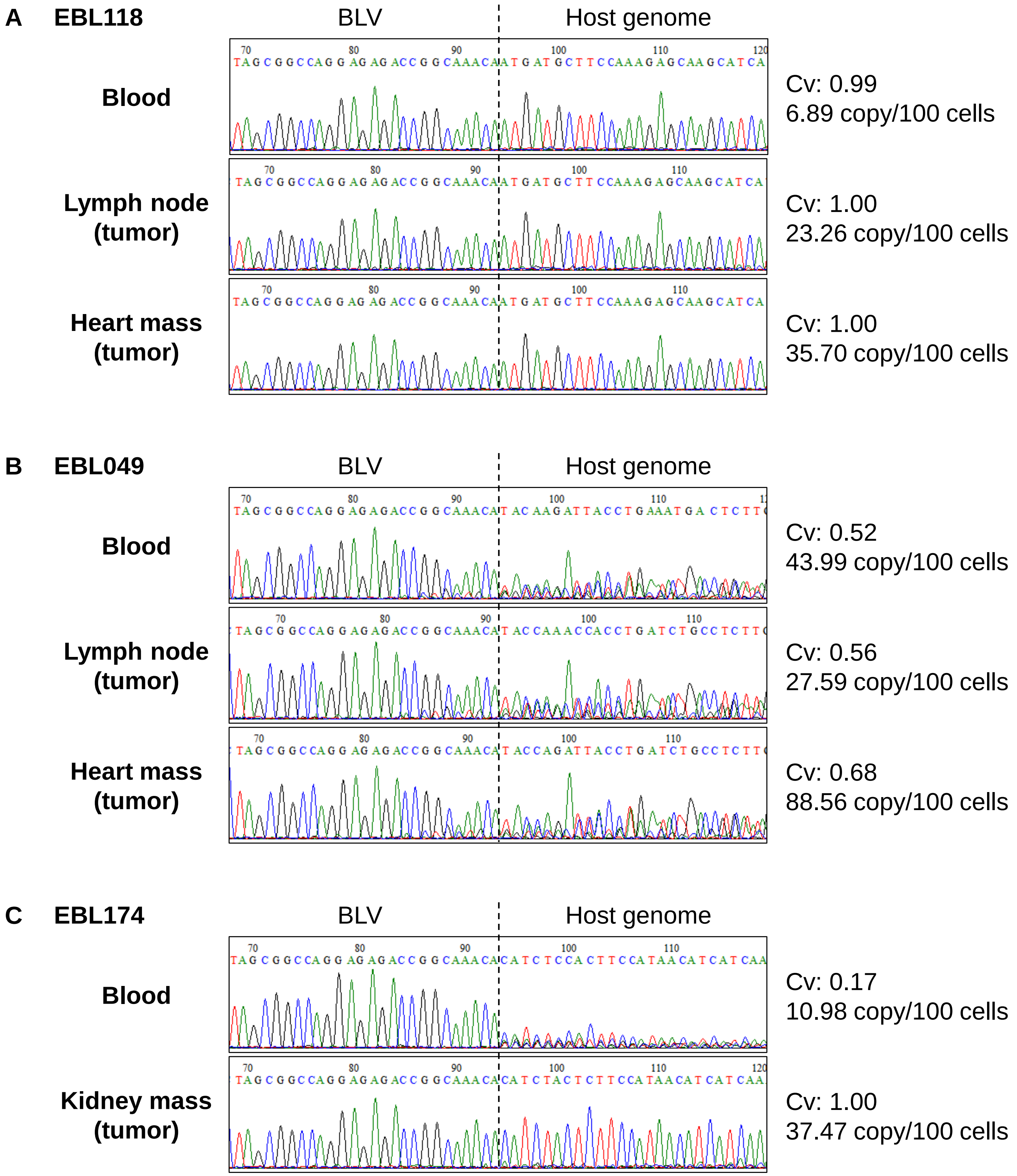


**Supplemental Fig. 2. Comparison of integration sites analyzed by RAISING-CLOVA in tumor and blood samples of EBL cattle.**

(A─C) Sangar sequence spectrum of representative blood and tumor samples from EBL cattle (A, EBL118; B, EBL049; C, EBL174) were shown with Cv and BLV PVL. Dashed line indicates the position of the BLV integration site.

#### Supplemental Table 1. Primers used in this study

| Primer name | Primer sequence | Length (bp) | Tm (°C) | GC (%) | Experiment | Reference |
|---|---|---|---|---|---|---|
| BLV-F1 | ATGAATGGCTCTCCCGCCTTTTT | 23 | 71.1 | 48 | ssDNA synthesis | Wada et al., 2022 |
| BLV-F2 | CTATCCGGCAGCGGTCAGGTAAG | 23 | 71.3 | 61 | dsDNA synthesis & 1st PCR (forward) | Wada et al., 2022 |
| NV-oligo-dT-ADP1 | ACAGCAGGTCAGTCAAGCAGTATTTTTTTTTTTTTTTTTTTTTTTVN | 47 | 75.0~77.1 | 23~28 | 1st PCR (reverse) | Wada et al., 2022 |
| BLV-F3 | ACACTCTTTCCCTACACGACGCTCTTCCGATCTCACCCGCGTTYGYTTCCTGTCTT | 54 (21) | 89.3 (70.4) | 54 (57) | 2nd PCR (forward) | Wada et al., 2022 |
| ADP1-HTS-R1 | GTGACTGGAGTTCAGACGTGTGCTCTTCCGATCTACAGCAGGTCAGTCAAGCAGTA | 56 (22) | 87.6 (63.9) | 52 (50) | 2nd PCR (reverse) | Wada et al., 2022 |
| Sequencing primer | ACACTCTTTCCCTACACGAC | 20 | 58.2 | 50 | Sangar sequencing | Illumina (San Diego, CA, USA) |

Adaptor sequence of each primer was shown in blue.

The length, Tm, and GC content of each primer without adaptor sequence were shown in brackets.

#### Supplemental Table 2. Reagents used in each step of RAISING

##### Step 1: ssDNA synthesis

| Components | Company | Volume (µL) |
|---|---|---|
| Genomic DNA (50 ng/µL) | - | 10.0 |
| 10× PCR Buffer for KOD-Plus-Neo | Toyobo | 5.0 |
| 2 mM dNTPs | Toyobo | 5.0 |
| 25 mM $MgSO_4$ | Toyobo | 3.0 |
| 10 µM F1 primer | Hokkaido System Science | 1.5 |
| KOD-Plus-Neo (1 U/µL) | Toyobo | 1.0 |
| Ultrapure water (DDW) | - | 24.5 |
| **Total** | | **50.0** |

##### Step 2: Column purification

ssDNA was purified using a Monarch PCR & DNA Cleanup Kit (New England Biolabs).
The purified ssDNA was eluted in 9.6 µL of DDW.

##### Step 3: Poly(AG)-tailing

###### 3-1. Poly(A)-tailing

| Components | Company | Volume (µL) |
|---|---|---|
| ssDNA | - | 8.2 |
| 10× TdT Buffer | New England Biolabs | 1.1 |
| 2.5 mM $CoCl_2$ | New England Biolabs | 1.1 |
| 10 mM dATP | New England Biolabs | 0.35 |
| TdT (20 U/µL) | New England Biolabs | 0.25 |
| **Total** | | **11.0** |

###### 3-2. Poly(G)-tailing

| Components | Company | Volume (µL) |
|---|---|---|
| PolyA-tailed ssDNA | - | 11.0 |
| 10× TdT Buffer | New England Biolabs | 0.1 |
| 2.5 mM $CoCl_2$ | New England Biolabs | 0.1 |
| 10 mM dGTP | New England Biolabs | 0.35 |
| DDW | - | 0.45 |
| **Total** | | **12.0** |

##### Step 4: dsDNA synthesis & 1st PCR

| Components | Company | Volume (µL) |
|---|---|---|
| PolyAG-tailed ssDNA | - | 12.0 |
| 5× Q5 Reaction Buffer | New England Biolabs | 12.0 |
| 10 mM dNTPs | Thermo Fisher Scientific | 1.2 |
| 10 µM F2 | Hokkaido System Science | 3.0 |
| 10 µM Oligo-dT-AD2 | Hokkaido System Science | 3.0 |
| Q5 HS-High-Fidelity DNA polymerase (2 U/µL) | New England Biolabs | 0.6 |
| DDW | - | 28.2 |
| **Total** | | **60.0** |

##### Step 5: 2nd PCR

| Components | Company | Volume (µL) |
|---|---|---|
| 1st PCR product (diluted 1:200 with DDW) | - | 1.0 |
| 10× PCR Buffer for KOD-Plus-Neo | Toyobo | 5.0 |
| 2 mM dNTPs | Toyobo | 5.0 |
| 25 mM $MgSO_4$ | Toyobo | 3.0 |
| 10 µM F3 primer | Hokkaido System Science | 1.5 |
| 10 µM ADP1-NGS-R1 | Hokkaido System Science | 1.5 |
| KOD-Plus-Neo (1 U/µL) | Toyobo | 1.0 |
| DDW | - | 32.0 |
| **Total** | | **50.0** |

#### Supplemental Table 3. Reaction conditions of each step of RAISING

##### Step 1: ssDNA synthesis

| Condition | Temperature | Time | Cycle |
|---|---|---|---|
| Pre-denaturation | 94°C | 2 min | 1 |
| Denaturation | 98°C | 10 sec | 25 |
| Annealing & extension | 68°C | 75 sec | |

##### Step 3: Poly(AG)-tailing

###### 3-1. Poly(A)-tailing

| Condition | Temperature | Time | Cycle |
|---|---|---|---|
| Poly(A)-tailing | 37°C | 30 min | |

###### 3-2. Poly(G)-tailing

| Condition | Temperature | Time | Cycle |
|---|---|---|---|
| Poly(G)-tailing | 37°C | 15 min | |
| Inactivation | 75°C | 10 min | |

##### Step 4: dsDNA synthesis & 1st PCR

| Condition | Temperature | Time | Cycle | |
|---|---|---|---|---|
| Pre-denaturation | 65°C | 5 min | 1 | dsDNA synthesis |
| Annealing | 64°C | 10 sec | 1 | |
| | 62°C | 10 sec | | |
| | 60°C | 10 sec | | |
| | 58°C | 10 sec | | |
| | 56°C | 10 sec | | |
| | 54°C | 10 sec | | |
| | 52°C | 10 sec | | |
| Extension | 72°C | 90 sec | | |
| Pre-denaturation | 94°C | 30 sec | 1 | 1st PCR |
| Denaturation | 98°C | 10 sec | 22 | |
| Annealing | 68°C | 10 sec | | |
| Extension | 72°C | 1 min | | |

##### Step 5: 2nd PCR

| Condition | Temperature | Time | Cycle |
|---|---|---|---|
| Pre-denaturation | 94°C | 2 min | 1 |
| Denaturation | 98°C | 10 sec | 30 |
| Annealing & extension | 68°C | 1 min | |
